# Contamination as a major factor in poor Illumina assembly of microbial isolate genomes

**DOI:** 10.1101/081885

**Authors:** Haeyoung Jeong, Jae-Goo Pan, Seung-Hwan Park

**Affiliations:** Infectious Disease Research Center, Korea Research Institute of Bioscience and Biotechnology (KRIBB), 125 Gwahak-ro, Yuseong-gu, Daejeon 34141, Republic of Korea; Biosystems and Bioengineering Program, University of Science and Technology (UST), 217 Gajeong-ro, Yuseong-gu, Daejeon 34113, Republic of Korea

## Abstract

The nonhybrid hierarchical assembly of PacBio long reads is becoming the most preferred method for obtaining genomes for microbial isolates. On the other hand, among massive numbers of Illumina sequencing reads produced, there is a slim chance of re-evaluating failed microbial genome assembly (high contig number, large total contig size, and/or the presence of low-depth contigs). We generated Illumina-type test datasets with various levels of sequencing error, pretreatment (trimming and error correction), repetitive sequences, contamination, and ploidy from both simulated and real sequencing data and applied k-mer abundance analysis to quickly detect possible diagnostic signatures of poor assemblies. Contamination was the only factor leading to poor assemblies for the test dataset derived from haploid microbial genomes, resulting in an extraordinary peak within low-frequency k-mer range. When thirteen Illumina sequencing reads of microbes belonging to genera *Bacillus* or *Paenibacillus* from a single multiplexed run were subjected to a k-mer abundance analysis, all three samples leading to poor assemblies showed peculiar patterns of contamination. Read depth distribution along the contig length indicated that all problematic assemblies suffered from too many contigs with low average read coverage, where 1% to 15% of total reads were mapped to low-coverage contigs. We found that subsampling or filtering out reads having rare k-mers could efficiently remove low-level contaminants and greatly improve the *de novo* assemblies. An analysis of 16S rRNA genes recruited from reads or contigs and the application of read classification tools originally designed for metagenome analyses can help identify the source of a contamination. The unexpected presence of proteobacterial reads across multiple samples, which had no relevance to our lab environment, implies that such prevalent contamination might have occurred after the DNA preparation step, probably at the place where sequencing service was provided.

## INTRODUCTION

During the last two decades, everyone witnessed how innovations in genome sequencing technologies have revolutionized almost all fields of biomedical research and application. Microbiology is one of the main beneficiaries among them, as next-generation genome sequencing has become the most cost-effective solution to answer questions of function, evolution, and interaction involving microbes and their surrounding environments (1). In the early years, massive amounts of low-quality short reads with data types totally different from what we have known have posed challenges to bioinformatics, especially for *de novo* assembly of genomes using Illumina sequencing platform (2‐4). Recent development and improvement in genome sequencing have resulted in the availability of several dozen software. We have to choose the most suitable one for a specific purpose. For example, there are more than forty *de novo* assemblers (https://en.wikipedia.org/wiki/Sequence_assembly). Best practice based on assembler evaluations (5, 6) or the use of automated assembly workbenches depending on validation or integration of outputs from multiple assemblers (7, 8) could help us obtain optimal Illumina assemblies of microbial isolate genomes.

The introduction of so-called “third generation sequencing” technologies (9) virtually monopolized by PacBio’s long-read SMRT^TM^ sequencing data and nonhybrid hierarchical genome assembly process (HGAP) (10) have greatly facilitated the completion of bacterial genomes using as few as one SMRT^TM^ cell which usually covers ~100x sequencing depth (11). This approach can dramatically reduce the labor and cost required for finishing genomes compared to first or second generation sequencing platforms. There are increasing cases of finishing bacterial genomes of interest using PacBio platform even in the presence of sequence reads that were produced using second generation sequencing technologies. However, the previous sequencing reads are not be usually utilized by HGAP anymore. This implies that more and more draft genome sequences, whether accurate or inaccurate, will have little chances of being re-evaluated if they are not going to be completed.

Carrying out hundreds of microbial sequencing project using Illumina platform as a usual customer of sequencing facilities, we encountered cases where *de novo* assembly resulted in too many contigs. Sometimes these assemblies comprised of short ones with low average coverages. Sometimes the total contig length was significantly larger than the estimated genome size despite sufficient sequencing depth. We learned from our experiences that pretreatment of reads, parameter scanning during the assembly procedure (mostly by changing k-mer length), subsampling reads, and using assemblers such as Velvet (12) that apply automatic coverage cutoff could improve our results for some problematic datasets. However, we did not take systematic approaches to find out the factors causing bad assemblies. Although there are many issues relevant to the inherent accuracy of large genome assemblies such as missing sequences, order/orientation errors, and erroneous reconstruction of repeats (13), these are beyond the scope of this study.

K-mer frequency analysis is an alignment-free method that is widely used in many aspects of genome researches, such as estimating genome sizes (14), diagnosing sequencing data quality and complexity (15), and pretreating massive sequencing reads (16). Theoretically, k-mer abundance in high-depth sequencing reads from an ideal haploid genome is distributed as a Gaussian with an average abundance equal to the sequencing depth. Low frequency k-mers represent Illumina-specific errors, genome heterogeneity, and contamination.

In this study, using test datasets and real Illumina sequence reads produced from a single run of 11 *Bacillus* and two *Paenibacillus* strains, we found extraordinary peak of rare k-mers from samples with failed assemblies due to contamination. We also demonstrated that contamination was the sole cause that spoiled genome assemblies in this case. Using tools were originally designed for metagenomics sequence analysis, the proportion of contamination in sequencing data and their possible source could be inferred. We were able to significantly improve the *de novo* assemblies of problematic samples by filtering out low abundant k-mers which efficiently eliminated low-level contamination from the reads.

## MATERIALS AND METHODS

### Construction of test datasets

Illumina reads that were produced from previous studies (17‐19), which are available at NCBI sequence read archive (SRA) under the accessions SRP058110 (*Escherichia coli* BL21), SRP058116 (*Bacillus subtilis* KCTC 1028), and SRP058417 (*Shigella boydii* ATCC 9210), were utilized to generate test dataset. 100-fold artificial Illumina reads were also simulated from the complete genome sequences of *E. coli* BL21 (CP010816.1), *Escherichia coli* K-12 MG1655 (NC_000913.3), *Saccharomyces cerevisiae* S288c (BK006934.2-BK006949.2, and AJ011856.1), and diploid yeast *Candida albicans* SC5314 (assembly 22, downloaded from http://www.candidagenome.org/) using ART (20) version 03-19-2015 with parameters -l 101 -ss HS25 -f 100 -m 400 -s 80. To simulate low-quality sequencing reads, error rate parameters of ART (-ir 0.00009 -ir2 0.00015 -dr 0.00011 -dr2 0.00023 for default setting) were multiplied appropriately. Unless otherwise mentioned, real Illumina reads were cut at 100x sequencing depth before being further processed.

### Bacterial strains, growth condition, and genome sequencing

Eleven *Bacillus* and two *Paenibacillus* strains chosen for genome sequencing are listed in Table 1. They are probiotic strains isolated from Korean traditional food (21), endophytic bacteria from cactus, and strains deliberately chosen from culture collection for biotechnological applications. Cells were grown aerobically in tryptic soy media at 30°C. Genomic DNA was isolated using Wizard Miniprep kit (Promega, Madison, Wisconsin, USA). Library was constructed using Truseq DNA sample prep kit. Paired end sequence reads (101-nt) were obtained using Illumina HiSeq 2000 platform (San Diego, California, USA) at National Instrumentation Center for Environment Management, Seoul National University (Seoul, Republic of Korea). Through sample multiplexing, sequencing reads from all thirteen libraries were obtained from a single run.

**Table 1.** List of bacterial strains and sequencing summary. (Pretreatments and *de novo* assembly were performed using CLC Genomics Workbench version 8.5. Trim rates based on the number of reads ranged from 80.24% to 92.02%).

| Sample ID | Strain description | Availability | No. of reads | No. of basepairs | No. of contigs<sup>a</sup> | Total contig length | N50 |
| --- | --- | --- | --- | --- | --- | --- | --- |
| Bpf | "*Bacillus polyfermenticus*" Bpf | Lab collection | 29,008,528 | 2,929,861,328 | 90 (63) | 4,127,902 | 502,253 |
| Bc1 | *Bacillus cereus* ATCC 4342 | ATCC | 38,787,810 | 3,917,568,810 | 6,052 (4,312) | 10,444,843 | 8,376 |
| Bc2 | *Bacillus cereus* ATCC 31382 | ATCC | 24,280,592 | 2,452,339,792 | 119 (56) | 9,286,527 | 443,015 |
| BaD11 | *Bacillus licheniformis* SCK B11 | T.-B. Um (21) | 31,066,026 | 3,137,668,626 | 84 (51) | 4,271,421 | 267,419 |
| Bm1 | *Bacillus megaterium* Bm1<sup>b</sup> | Y. Bashan | 26,594,682 | 2,686,062,882 | 133 (93) | 5,279,131 | 1,118,798 |
| Bm2 | *Bacillus megaterium* Bm2<sup>b</sup> | Y. Bashan | 26,524,590 | 2,678,983,590 | 78 (52) | 5,113,416 | 1,121,613 |
| Bp1 | *Bacillus pumilus* Bp1<sup>b</sup> | Y. Bashan | 43,302,480 | 4,373,550,480 | 3,638 (2,472) | 7,123,372 | 61,549 |
| Bp2 | *Bacillus pumilus* Bp2<sup>b</sup> | Y. Bashan | 39,622,506 | 4,001,873,106 | 44 (33) | 3,683,921 | 968,875 |
| ES4 | *Bacillus pumilus* ES4<sup>b</sup> | Y. Bashan | 29,157,078 | 2,944,864,878 | 69 (57) | 3,695,884 | 969,001 |
| Rizo | *Bacillus pumilus* Rizo<sup>b</sup> | Y. Bashan | 39,163,182 | 3,955,481,382 | 191 (179) | 3,743,083 | 968,988 |
| Bsu | *Bacillus subtilis* Bsu<sup>b</sup> | Y. Bashan | 23,086,814 | 2,331,768,214 | 64 (51) | 4,078,262 | 1,055,276 |
| Pan | *Paenibacillus antarticus* KCTC 13016 | KCTC | 24,832,632 | 2,508,095,832 | 129 (95) | 5,373,088 | 385,650 |
| Pgl | *Paenibacillus glacialis* KCTC 13874 | KCTC | 41,997,612 | 4,241,758,812 | 177 (97) | 5,714,841 | 217,285 |

<sup>a</sup> Numbers in parentheses designate the number of contigs shorter than 1,000 bp.
<sup>b</sup> Endophytic bacteria isolated from cacti.

### Pretreatment of sequence reads and *de novo* assembly

Trimmomatic (22) included in the A5-Miseq package (23) version 20140604 was used for adaptor sequence removal and quality trimming using modified parameters (SLIDINGWINDOW:4:20 MINLEN:75) to obtain longer sequences. SGA (24) version 0.10.13 was used for k-mer-based error correction when required. For k-mer abundance analysis, Jellyfish (14) version 2.2.3 or khmer (16) version 2.0 was used with a k-mer size of 20. Jellyfish, a k-mer counting program without filtering function, runs much faster than khmer but gives similar results. Therefore, Jellyfish was used for the initial screen while khmer was used to filter out k-mers below a specified coverage of problematic reads or for counting, format conversion, and general manipulation of paired fastq files such as interleaving and splitting. CLC Genomics Workbench versions 8.5 or 9.0 was used for *de novo* assembly with a word size of 64. Trimming (low quality limit 0.01, max 1 ambiguous nucleotide allowed per read, min length 50 nucleotides) was applied in case trimmomatic was skipped. Subsampling prior to assembly was also carried out in the same environment (i.e., CLC Genomics Workbench). Average read coverages of contigs were obtained by selecting mapping options from the *de novo* assembly tool in CLC Genomics Workbench. Contig comparison and visualization of assembly metrics were carried out using QUAST (25) version 2.3. Prokka (26) version 1.11 was then run for functional annotation of assembled genome sequences. Nucleotide and amino acid sequences of predicted genes were analyzed using specI (27) to identify species.

### Identification of 16S rRNA genes

Pairs of read files without pretreatment were converted to FASTA format and passed to REAGO (28) version 1.1 to retrieve reads originating from 16S rRNA genes and to reconstruct them separately (either from multiple copies of genes within a single strain or from contaminating genomes). The reconstructed 16S rRNA genes from REAGO (full genes only) and Prokka (full and partial genes) were analyzed using EzTaxon server (29).

### Estimation of contamination using metagenome analysis tools

To estimate the proportion of contaminated reads and identify the source organism at various phylogenetic levels, a couple of tools for metagenomics sequence analysis were applied. Raw reads without pretreatment were subjected to analysis with MetaPhyler (30) SRV0.115, an optimized version suitable for short reads, and to Kraken (31) version 0.10.5-beta using MiniKraken 20141208 as the reference genome database. They were also subjected to analysis with PhyloSift (32) version 1.0.0_01. Results were visualized using Krona (33). Kraken differs from the other two software in that it assigns taxonomic labels to each reads, whereas MetaPhyler and PhyloSift generate taxonomic profiles and phylogenetic analysis, respectively.

## RESULTS

### K-mer abundance profiles of test datasets

We prepared test dataset with various conditions that might affect *de novo* assemblies. We also counted their k-mer frequencies. Simulated reads with increased error rate showed much divergence at low k-mer frequencies (Fig. 1A). There was a combinatorial effect of sequential quality trimming and error correction of reads on k-mer distribution at low frequencies with data volume adjustment so that all treated reads could have the same amounts of final basepairs (Fig. 1B). Interestingly, error rate and pretreatment did not appear to be the major factor affecting the assembly metrics if adequate sequencing coverage was ensured without contamination (Supplementary Table S1). Real sequencing reads from repeat-rich genome *Shigella boydii* produced as many as 443 contigs (4.53 Mb) due to the presence of hundreds of copies of insertion sequences which resulted in highly fragmentary assembly with a shoulder peak just beyond the main one, representing k-mers from repetitive sequences (Fig. 1C). Although *E. coli* BL21 also showed multiple peaks at high frequency k-mer range, the relative height of the highest secondary peak to the main was only 0.23%. It was 45.3% for *Shigella boydii.*

**Fig. 1.**
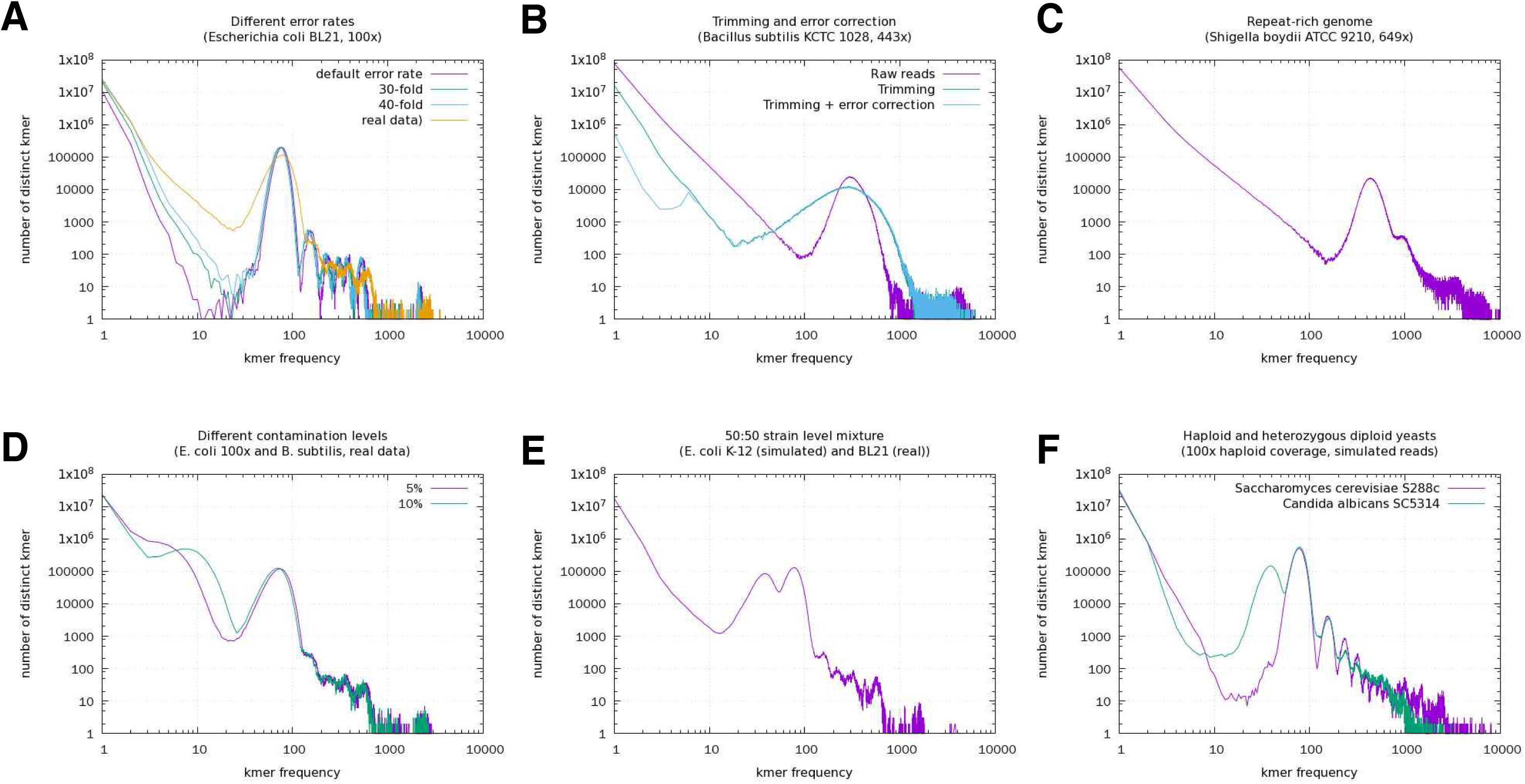
Abundance profiles of 20-mers in test dataset. Sequencing depth of all dataset are 100x unless otherwise mentioned. All axes are shown in log scale. A, Simulated and real reads from *E. coli* BL21 with different errors. B, Real reads from *B. subtilis* KCTC 1028 with either trimming or error correction. All reads were adjusted to 443x (final dataset). C, Real reads at 640x from repeat-rich genome of *Shigella boydii* ATCC 9210. D*, E. coli* BL21 reads contaminated with 5% or 10% *B. subtilis* KCTC 1028 reads. E, 50:50 mixture of simulated reads from *E. coli* K-12 MG1655 and real reads from *E. coli* BL21. F, Simulated reads from diploid yeast *Saccharomyces cerevisiae* S288c and heterozygous diploid yeast *Candida albicans* SC5314.

Contamination appeared to be the major factor that deteriorated both assembly results and k-mer profiles (Fig. 1D). There was a secondary peak in the low frequency range with different peak locations and heights. We also carried out *de nov*o assemblies with various contamination levels. Compared to species level contamination where 5-10% contamination mostly spoiled the assemblies, strain level contamination resulted in the worst assembly when 50% mixture was used (Supplementary Table S1 and Fig. S1). For the latter situation, k-mer spectrum is shown in Fig. 1E. It is reminiscent of a *de novo* assembly of a heterozygous diploid genome. We also compared k-mer spectra of simulated reads of haploid yeast *Saccharomyces cerevisiae* S288c and heterozygous diploid yeast *Candida albicans* SC5314 (Fig. 1F). Reads from haploid genome showed a major peak around sequencing depth (100x) and minor peaks arising from repeats at high frequencies. On the contrary, SC5314 showed the largest secondary peak left to the main peak. As expected, heterozygous diploid genome resulted in much worse assembly that haploid genome (Supplementary Table S1).

### Bad assemblies from real data were associated with anomalous k-mer abundant profile

Real Illumina sequencing data produced from one single multiplexed run of thirteen bacterial isolate genomes were preprocessed (trimming only) and assembled using CLC Genomics Assembler. At first glance, samples Bc1 and Bp1 failed in *de novo* assembly as they showed extraordinarily high contig numbers. In addition, the total contig lengths were much larger than the expected genome size (Table 1). Although Sample Bc2 resulted in a moderately large contig number that might be regarded as normal, it should be regarded as a failure due to total contig length of nearly 9.3 Mb, which is impossible for a *Bacillus cereus* strain. All reads were pretreated using trimmomatic (adaptor removal and trimming) and subjected to Jellyfish analysis for k-mer counting. As expected, samples Bc1, Bc2, and Bp1 all showed a secondary peak at the rare k-mer range (Fig. 2).

**Fig. 2.**
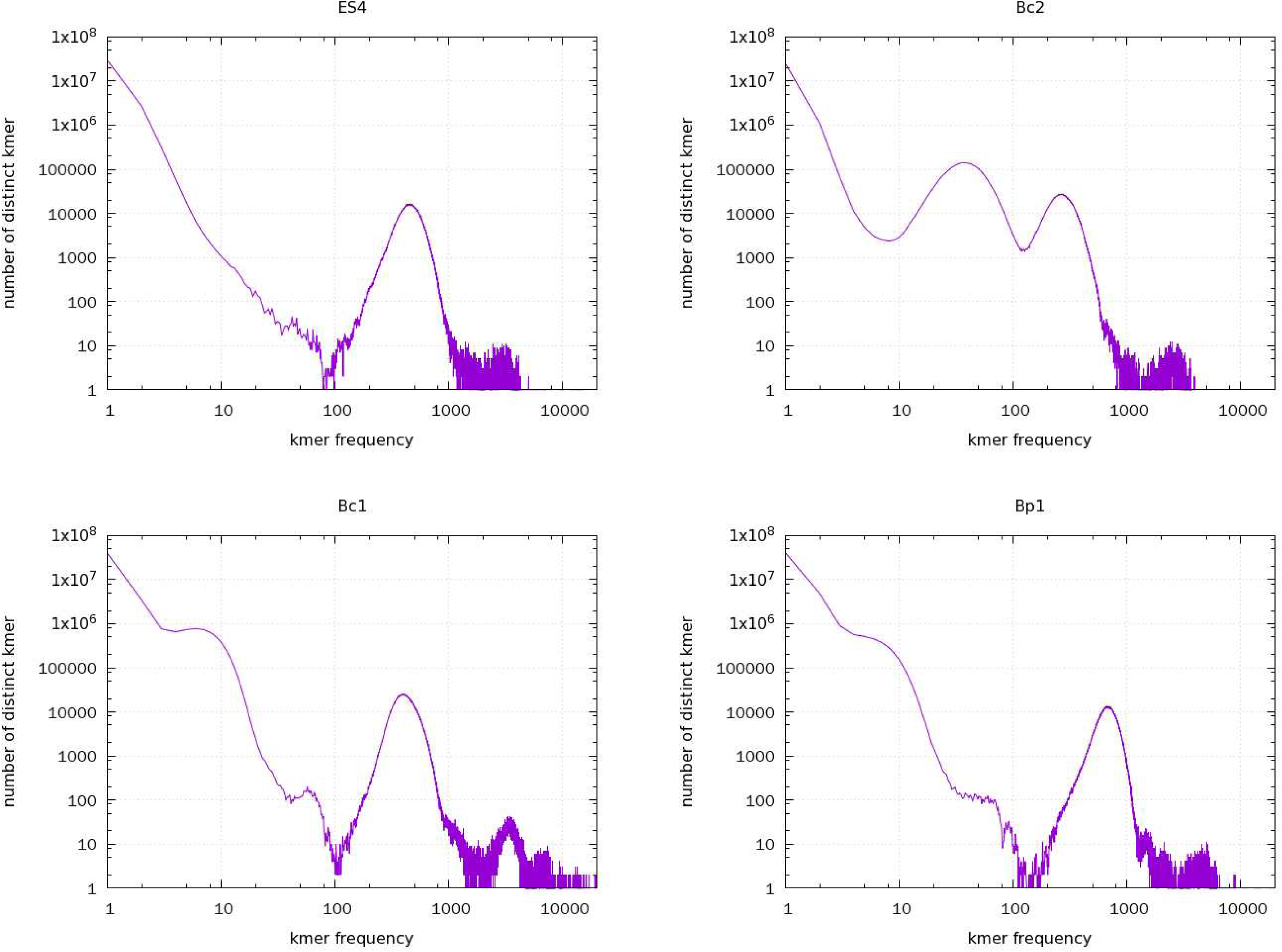
Abundance profiles of 20-mers in selected sequence reads.

### Sample Bp1 was contaminated by a GC-rich genome

While inspecting QUAST report for graphical representation of assembly results, we found that Bp1 contained contigs whose %GC was much higher (~70% on average) than what was expected for a *Bacillus pumilus* genome (Supplementary Fig. S2A). Contigs could be divided at 55% G+C into a low-GC group (172 contigs, total length of 3,759,031 bp, N50 of 906,842 bp) and a high-GC group (3,466 contigs, total length of 3,364,341 bp, N50 of 1,280 bp). BLAST analysis of the largest contig (17,311 bp) of the second group revealed that the high GC contigs originated from GC-rich *Caulobacter* species. GC-rich contamination in sample Bp1 was not conspicuous at the read level (Supplementary Fig. 2B) because only 2.91% of total reads were ≥ 55% G+C.

### Read subsampling or removing low-abundant k-mers improved *de novo* assemblies

As a fast measure to improve the problem in assembly, reads were subsampled at 5-50% ranges and subject to *de novo* assembly. 5% of subsample yielded far better assemblies than using the whole dataset for Bc1 and Bp1 (Fig. 3). Based on k-mer spectrum analysis, the atypical peak at low k-mer frequencies disappeared gradually with decreasing subsample size. The shortcoming of this treatment is the difficulty in choosing the proper sample size to abolish most reads containing low-abundant k-mers (probably due to contamination) while keeping adequate sequencing depth. When the total contig size calculated from the assembly of filtered reads, a better estimate for genome size, was used for the calculation of sequencing depth of subsamples, 5% of reads corresponded to 31.7x (Bc2) or 48.7x (Bp1) coverages. These are slightly under the optimal level to ensure successful assemblies. A still better solution was to perform assembly after read filtering under a specified k-mer coverage (50 for Bc1 and Bp1; 100 for Bc1). This produced as little as 30 contigs. However, Bc2 assembly was not improved using subsampling or k-mer filtering because the two k-mer peaks significantly overlapped with each other.

**Fig. 3.**
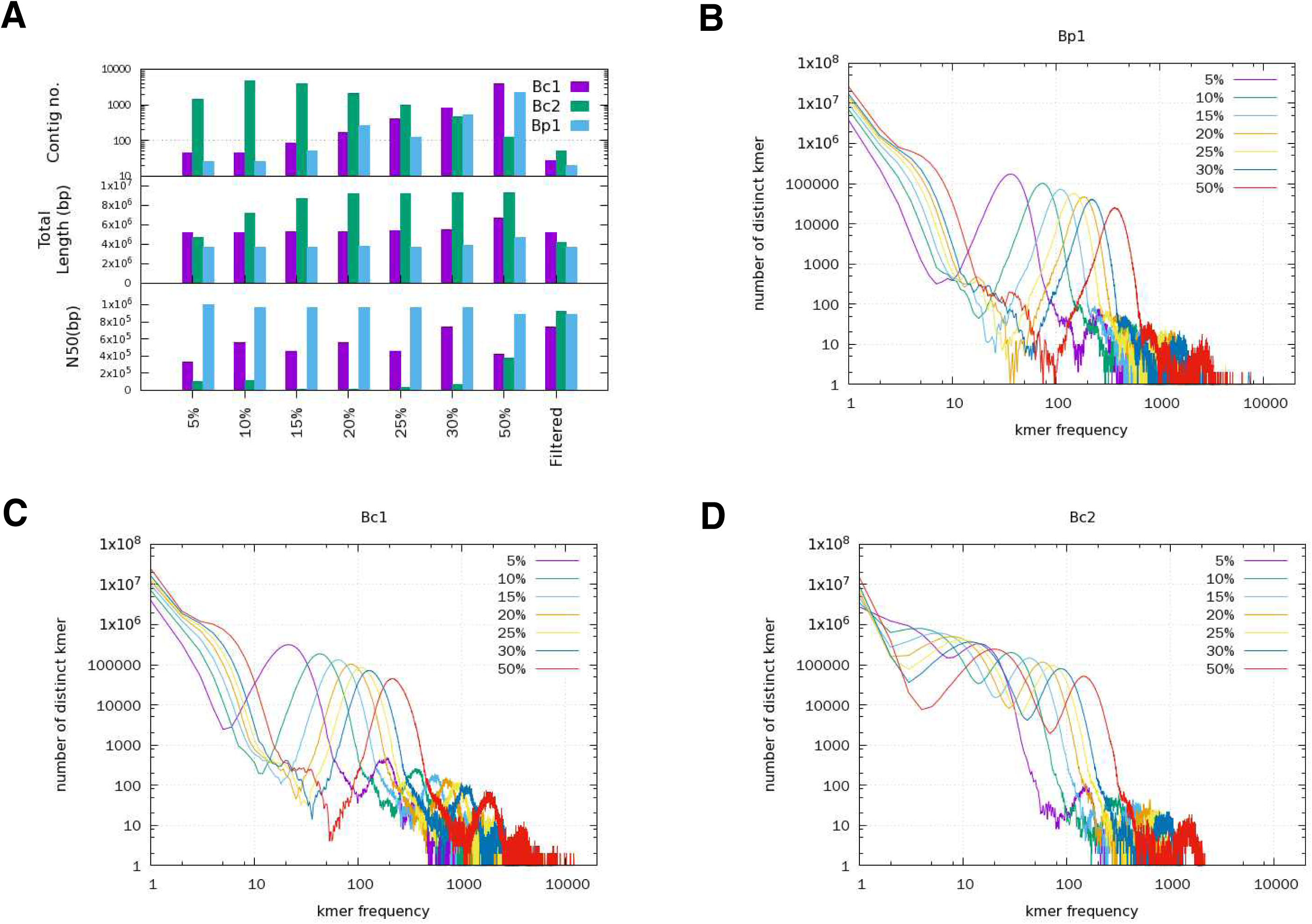
Effect of subsamplig or filtration on *de novo* assembly. A, Comparison of assembly results. B-D, Changes in k-mer distribution of reads caused by subsampling for Bp1, Bc1, or Bc1.

### High contig numbers of failed assemblies were due to short and low-coverage contigs

Cumulative contig length reached a plateau as short contigs were added (Fig. 4A), implying that longer contigs contributed the most to the total length. However, in failed assemblies with extraordinarily high contig numbers such as Bc1 and Bp1, short contigs accounted for half of the total contig length (Fig. 4B). Next, we plotted average read coverage, length, and cumulative length of contigs sorted in the increasing order of read coverage (Fig. 5). Normal assemblies showed similar pattern to Bp2, where an average read coverage leap (designated by two dashed lines) divided contigs into two groups. High coverage contigs contained sequences present once or more (repeats) that could be expected at a given sequencing depth. Contigs belonging to the low coverage group were very short (mostly < 1 kb). They contributed less to the total contig length in normal assemblies. For Bc1 and Bp1, short contigs with low coverage accounted for the majority of contig numbers. Quite high numbers of reads were mapped to low coverage contigs for poor assemblies, such as 1.5% for Bc1, 15.0% for Bc2, and 1.1% for Bp1. In normal assemblies, only 1,132 – 5,696 reads out of tens of million reads were mapped to low coverage contigs (Supplementary Table S2). Total length of low coverage contigs were also very long (3.45 – 5.21 Mb) for sample Bc1, Bc2, and Bp1. It was noteworthy that Bc2 had contigs with mid-level coverage. Their lengths were comparable to those in the high coverage group (Fig. 5). In contrast to Bc1 and Bp1, the low-to-mid level coverage group of Bc2 could be regarded as a high level of assembly (Supplementary Table S3). However, we could not designate any single species to it because neither 16S rRNA gene sequence nor definitive specI analysis result (78.11% average identity to *Bacillus* sp. 2_A_57_CT2, GenBank ACWD01000000) was obtained. We also found that the *de novo* assembly of artificially contaminated test dataset of 100x Illumina reads from *E. coli* BL21 containing 5% *B. subtilis* ATCC 1028 reads showed similar pattern of length-coverage distribution as Bc1 and Bp1 (Supplementary Fig. S4). Out of 4,577 contigs, 4,475 contigs with total length of 1,327,810 bp had low coverage (under 50x). Re-mapping after assembly revealed that low coverage contigs had only 89,988 reads (1.9% of total reads), where 89,594 reads of them were derived from *B. subtilis* genome. This observation further supports that short and low-coverage contigs in failed assembly are due to contaminating reads.

**Fig. 4.**
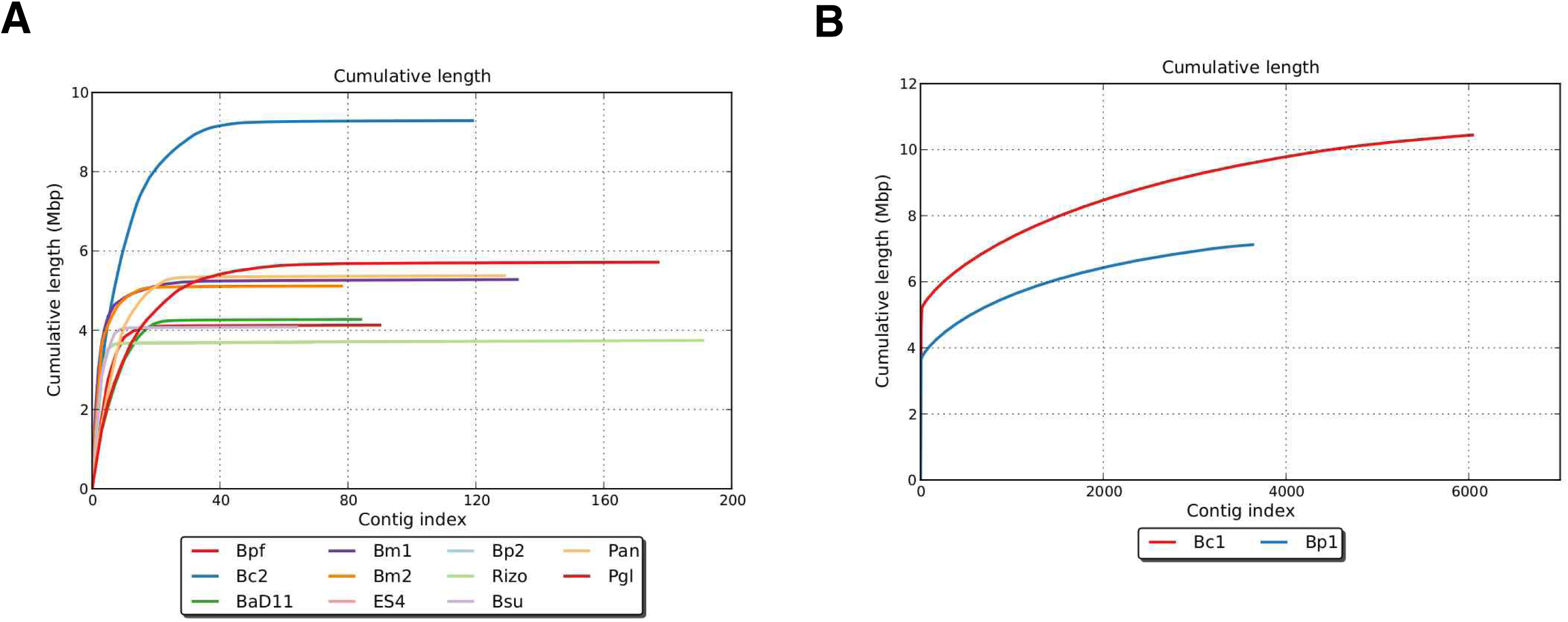
Cumulative length plot of reversely sorted contigs. A, Eleven of the 13 samples (except Bc1 and Bp1). B, Plot for Bc1 and Bp1.

**Fig. 5.**
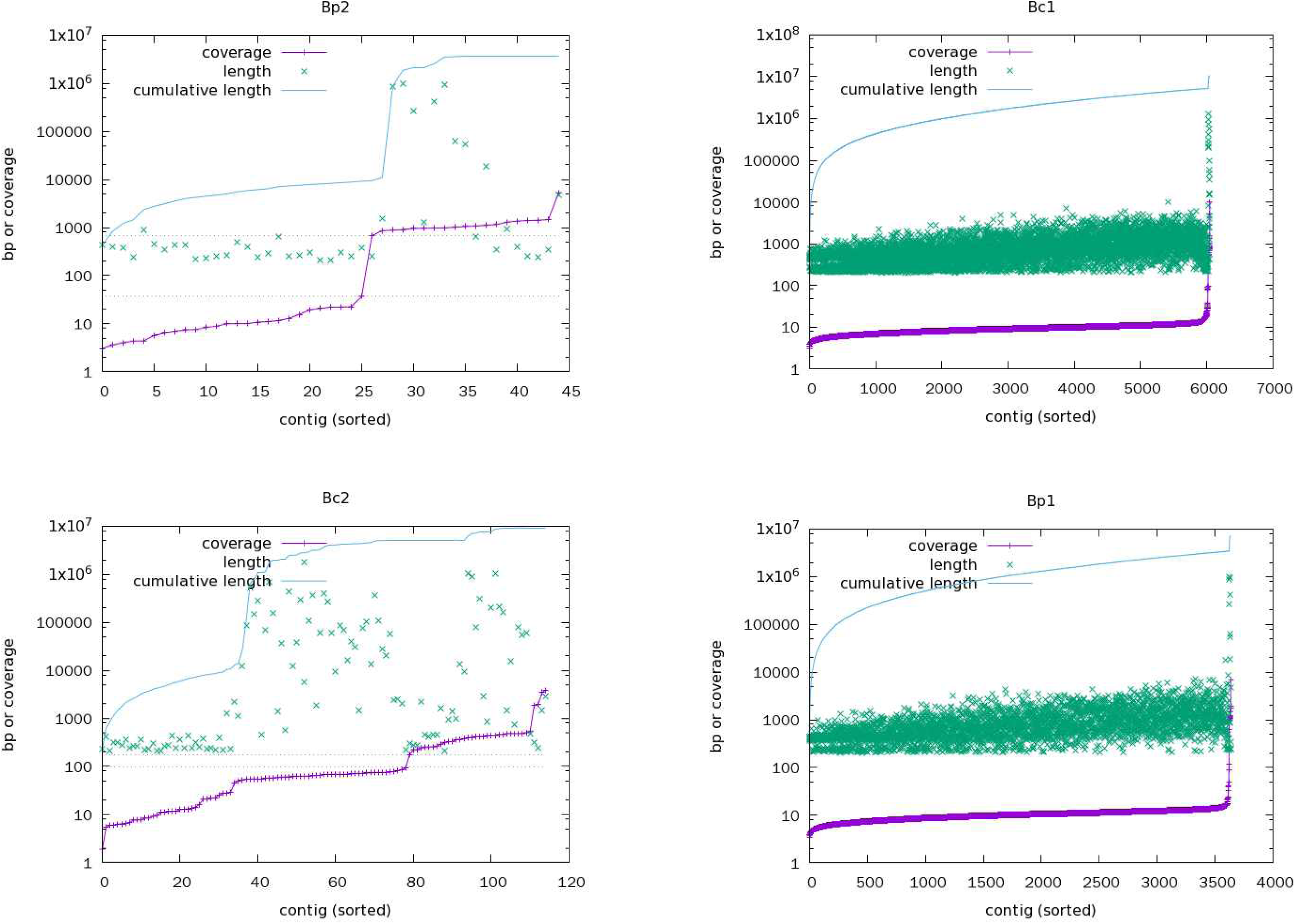
Coverage-length plot of assemblies. Contigs are sorted in increasing order.

### Identification of multiple 16S rRNA genes from reads and contigs

Because it became apparent that contamination in sequence reads was the main factor causing a large number of small contigs with low coverage, eventually leading to spoiled assembly, we determined the identities of contaminant. When read files without pretreatment were analyzed using REAGO (requiring reads of a fixed length), multiple full-length 16S ribosomal RNA genes were found from eleven out of thirteen samples. Because REAGO could distinguish ribosomal gene copies from a single genome, all full-length 16S ribosomal RNA genes were searched against database using EzTaxon server. 16S rRNA genes either full or partial were also identified from Prokka annotation of assembled sequences, where 16S rRNA genes from a single species were normally found from overcollapsed contigs. Results are summarized in Table 2. We could not find contaminating 16S rRNA genes in full form from REAGO results. However, fragmented caulobacterial sequences were found from at least five samples.

**Table 2.** Identification of 16S rRNA genes from reads or assemblies and their EzTaxon analysis results. (Copies of 16S rRNA genes identified by AGO for samples Bpf, Bc2, and Bsu showed different top hits due to intragenomic variations. In such cases, results that were the most similar to Prokka e chosen)

| Sample ID | From REAGO results<sup>a</sup> | From Prokka results |
| --- | --- | --- |
| Bpf | (2) *Bacillus siamensis* KCTC 13613(T) 99.87% | *Bacillus siamensis* KCTC 13613(T) 99.93% |
| Bc1 | Not identified | *Bacillus anthracis* ATCC 14758(T) 100%<br>(partial) *Paenibacillus glacialis* KGC91(T) 99.75% |
| Bc2 | (6) *Bacillus subtilis* subsp. *inaquosorum* KCTC 13429(T) 99.93% | (partial) *Bacillus subtilis* subsp. *inaquosorum* KCTC 13429(T) 100% |
| BaD11 | (7) *Bacillus licheniformis* ATCC 14580(T) 99.87%<sup>b</sup> | *Bacillus licheniformis* ATCC 14580(T) 99.93% |
| Bm1 | (10) *Bacillus megaterium* NBRC 15308 99.74%<sup>b</sup> | *Bacillus aryabhattai* B8W22(T) 100%<sup>c</sup> |
| Bm2 | (5) *Bacillus aryabhattai* B8W22(T) 99.87%<sup>b</sup> | *Bacillus aryabhattai* B8W22(T) 100% |
| Bp1 | (2) *Bacillus safensis* FO-36b(T) 99.93% | *Bacillus safensis* FO-36b(T) 100%<br>*Caulobacter mirabilis* FWC38(T) 97.39 |
| ES4 | (2) *Bacillus safensis* FO-36b(T) 100%<sup>b</sup> | *Bacillus safensis* FO-36b(T) 100% |
| Bp2 | (2) *Bacillus safensis* FO-36b(T) 99.87%<sup>b</sup> | *Bacillus safensis* FO-37b(T) 100% |
| Rizo | Not identified | *Bacillus safensis* FO-36b(T) 100% |
| Bsu | (4) *Bacillus subtilis* subsp. *inaquosorum* KCTC 13429(T) 99.87% | *Bacillus subtilis* subsp. *inaquosorum* KCTC 13429(T) 99.93% |
| Pan | (2) *Paenibacillus antarcticus* LMG 22078(T) 100% -all | *Paenibacillus antarcticus* LMG 22078(T) 100% |
| Pgl | (8) *Paenibacillus glacialis* KFC91(T) 99.60% | *Paenibacillus glacialis* KFC91(T) 99.65% |

<sup>a</sup>Numbers in parentheses indicate the number of full 16S rRNA gene identified using REAGO.
<sup>b</sup>Fragments of *Caulobacter mirabilis* FWC38(T) 16S rRNA genes were found.
<sup>c</sup>*Bacillus megaterium* NBRC 15308 (99.87%) ranked the second

Secondary 16S rRNA genes predicted by Prokka were found to belong to *Paenibacillus galicialis* and *Caulobacter mirabilis* (in Bc1 and Bp1, respectively). The presence of *Panibacillus glacialis* gene in sample Bc1 might reflect cross-contamination from sample Pgl. It should be noted that multiple samples, at least six, contained 16S rRNA gene sequence (partial or full) of *Caulobacter* species. However, no secondary 16S rRNA could be found from Bc2 using REAGO or Prokka, despite the completeness of contigs originating from contamination.

### Use of metagenome analysis tools suggests that caulobacterial contamination might be prevalent across samples

We applied MetaPhyler to all raw reads of the 13 samples to estimate the relative abundance of taxonomic units to rank phylum through genus. Results were compared at family level, the highest level where possible contamination across all samples could be maximally distinguishable (Table 3). Family *Caulobacteraceae* was detected from nine samples, including all six samples containing caulobacterial 16S rRNA genes from prior analysis. The abundance of *Caulobacteraceae* was the highest in Bp1 (0.46%) (Supplementary Fig. S5 for Krona plot). In Bc1, the abundance of *Paenibacillaceae* was 0.07%. In Bc2, the abundances of *Listeriaceae* (order *Bacillales*) and *Streptococcaceae* (order *Lactobacillaceae*) were 0.16% and 0.04%, respectively. Unexpectedly, in samples Bm1 and Bm2, the abundance of *Staphylococcaceae* and *Listeriaceae* (all order *Bacillaes*) was about 0.2%. It was also notable that Pan and Pgl belonging to genus *Paenibacillus* contained a significant amount of *Bacillus* reads (relative abundance of 6.44% and 3.55%, respectively).

**Table 3.** Results obtained from 13 sequence reads using metagenome analysis tools. (Results shown at family level, which was the highest taxonomy level that could difference between samples. Asterisks represent samples that contain caulobacterial 16S rRNA genes. Numbers given by PhyloSift are summed taxonomic mass heir ratios are available from Krona plot)

|  | Bpf | Bc1 | Bc2 | BaD11* | Bm1* | Bm2* | ES4* | Bp1* | Bp2* | Bsu | Pan | Pgl | Rizo |
| --- | --- | --- | --- | --- | --- | --- | --- | --- | --- | --- | --- | --- | --- |
| **Summary** |  |  |  |  |  |  |  |  |  |  |  |  |  |
| Laboratory | A | A | A | B | B | B | B | B | B | B | A | A | B |
| Total number of reads | 29,008,528 | 38,787,810 | 24,280,592 | 31,066,026 | 26,594,682 | 26,524,590 | 29,157,078 | 43,302,480 | 39,622,506 | 23,086,814 | 24,832,632 | 41,997,612 | 39,163,182 |
| MetaPhyler (# reads) | 64,659 | 77,771 | 64,661 | 77,552 | 34,915 | 31,742 | 79,104 | 79,216 | 75,009 | 83,433 | 4,091 | 4,994 | 83,554 |
| Kraken (# reads) | 22,812,318 | 29,763,422 | 19,231,872 | 26,189,384 | 20,097,293 | 16,433,313 | 6,045,238 | 8,925,429 | 8,184,870 | 19,858,356 | 1,257,652 | 545,962 | 8,089,394 |
| PhyloSift (probability) | 104,418 | 173,893 | 109,780 | 155,672 | 122,118 | 126,832 | 154,933 | 231,923 | 201,438 | 124,361 | 70,090 | 135,006 | 209,387 |
| Metaphyler (% classified) | 0.22% | 0.20% | 0.27% | 0.25% | 0.13% | 0.12% | 0.27% | 0.18% | 0.19% | 0.36% | 0.02% | 0.01% | 0.21% |
| Kraken (% classified) | 78.64% | 76.73% | 79.21% | 84.30% | 75.57% | 61.96% | 20.73% | 20.61% | 20.66% | 86.02% | 5.06% | 1.30% | 20.66% |
| PhyloSift (%) | 0.36% | 0.45% | 0.45% | 0.50% | 0.46% | 0.48% | 0.53% | 0.54% | 0.51% | 0.54% | 0.28% | 0.32% | 0.53% |
| **MetaPhyler<sup>c</sup>** |  |  |  |  |  |  |  |  |  |  |  |  |  |
| *Bacillaceae* | 99.52% | 99.88% | 99.25% | 99.67% | 97.53% | 97.74% | 99.81% | 99.38% | 99.82% | 99.47% | 6.44% | 3.55% | 99.81% |
| *Paenibacillaceae* | 0.01% | 0.07% |  |  |  |  |  |  |  |  | 91.66% | 94.47% | 0.00% (3)<sup>a</sup> |
| *Staphylococcaceae* |  |  |  |  | 0.15% | 0.16% |  |  |  |  |  |  |  |
| *Listeriaceae* |  |  | 0.16% |  | 0.04% | 0.04% |  |  |  |  |  |  |  |
| *Streptococcaceae* |  |  | 0.04% |  |  |  |  |  |  |  |  |  |  |
| *Proteobacteria* (phylum) |  |  | 0.01% | 0.07% | 0.23% | 0.1% | 0.09% | 0.56% | 0.04% | 0.06% |  |  | 0.07% |
| *Caulobacteraceae* | 0.00% (2)<sup>a</sup> |  |  | 0.03% | 0.17% | 0.07% | 0.04% | 0.46% | 0.03% | 0.03% |  |  | 0.04% |
| **Kraken<sup>c</sup>** |  |  |  |  |  |  |  |  |  |  |  |  |  |
| *Bacillaceae* | 94.28% | 99% | 95.74% | 99.35% | 98.96% | 99% | 97.03% | 96.35% | 97.27% | 99.2% | 6.16% | 14.57% | 97.01% |
| *Paenibacillaceae* |  | 0.01% |  |  | 0.02% |  | 0.07% | 0.08% | 0.07% |  | 9.49% | 49.32% | 0.07% |
| *Staphylococcaceae* |  |  | 0.01% |  |  | 0.01% | 0.02% | 0.02% | 0.02% |  |  |  | 0.02% |
| *Listeriaceae* |  |  | 0.01% |  |  |  | 0.05% | 0.05% | 0.05% |  |  | 0.31% | 0.05% |
| *Streptococcaceae* |  |  |  |  |  |  | 0.02% | 0.02% | 0.02% |  |  | 0.82% | 0.02% |
| *Proteobacteria* (phylum) | 5.01% | 0.04% | 0.16% | 0.04% | 0.08% | 0.04% | 0.41% | 1.01% | 0.37% | 0.03% | 75.56% | 2.01% | 0.35% |
| *Alcaligenaceae* | 0.82% | 0.01% | 0.01% |  | 0.02% |  | 0.04% | 0.15% | 0.05% |  | 14.99% | 0.27% | 0.02% |
| *Alteromonadaceae* | 4.18% | 0.02% | 0.14% | 0.01% | 0.02% | 0.01% | 0.2% | 0.27% | 0.2% | 0.01% | 61.36% | 0.92% | 0.14% |
| *Burkholderiaceae* |  |  |  |  | 0.01% | 0.01% | 0.02% | 0.08% | 0.01% |  |  |  | 0.02% |
| *Caulobacteraceae* |  |  |  | 0.01% | 0.02% | 0.01% | 0.04% | 0.23% | 0.02% | 0.01% |  |  | 0.05% |
| **PhyloSift<sup>c</sup>** |  |  |  |  |  |  |  |  |  |  |  |  |  |
| *Bacillaceae* | 94.90% | 95.79% | 89.64% | 95.96% | 94.22% | 94.41% | 94.84% | 93.81% | 95.05% | 95.63% | 2.24% | 1.33% | 94.98% |
| *Paenibacillaceae* |  | 1.52% |  |  |  |  |  |  |  |  | 97.81% | 97.97% |  |
| *Streptococcaceae* |  |  | 5.74% |  |  |  |  |  |  |  |  |  |  |
| *Caulobacterales* (order<sup>b</sup>) |  |  |  |  |  |  |  | 1.15% |  |  |  |  |  |

<sup>a</sup> Numbers in the parentheses indicate classified reads.
<sup>b</sup> Result at family level was not available.
5 ° Percentages represent relative abundance (MetaPhyler), ratio of classified reads (Kraken), and ratio of probability mass distribution (PhyloSift).

We then used Kraken to attach taxonomic label to each read after classification, where the resultant figures cannot be directly comparable with those obtained through MataPhyler or PhyloSift. Read classification rates across samples were significantly different in Kraken results. For examples, the classification rate was as high as 86.0% for Bsu reads. It was only 5.1% for Pan or 1.3% for Pgl. Because Kraken analysis depends on short exact alignments of reads on prebuilt genome library, its results are largely affected by the comprehensiveness of the library. Kraken reported much diverse and detailed phylogenetic distribution (due to possible contamination) than MetaPhyler (Supplementary Fig. S6 for Bp1). However, the actual percentage for reads classified to irrelevant taxons was below 0.1%, making it difficult to discriminate real contamination from noise that might be a function of reference database. In particular, proteobacterial reads appeared to be prevalent among all samples, where *Alcaligenaceae* (*Achromobacter xylosoxidans*) and *Altermonadaceae* (*Alteromonas mediterranea*) were the most common with a fairly similar proportion (Supplementary Fig. S7 for Bpf and Pan). Similar patterns were frequently found from Kraken analyses of Illumina reads derived from other totally unrelated sequencing projects (the ratio of *Achromobacter xylosoxidans* to *Alteromonas mediterranea* was approximately 1:1~10:1, data not shown). Therefore, similar proteobacterial distribution patterns across multiple samples (except for caulobacterial contamination) without 16S rRNA evidence were likely to be false positives.

Finally, all reads were subjected to PhyloSift analysis, a phylogeny-driven statistical hypothesis test to determine the presence of phylogenetic lineages where abundance information is given as probability distribution. PhyloSift showed the most conservative results among the three analysis tools (i.e., MetaPhyler, Kraken, and PhyloSift). PhyloSift analysis results revealed that contaminations occurred only in Bc1, Bc2, Bp1, Pan and Pgl (Fig. 6).

**Fig. 6.**
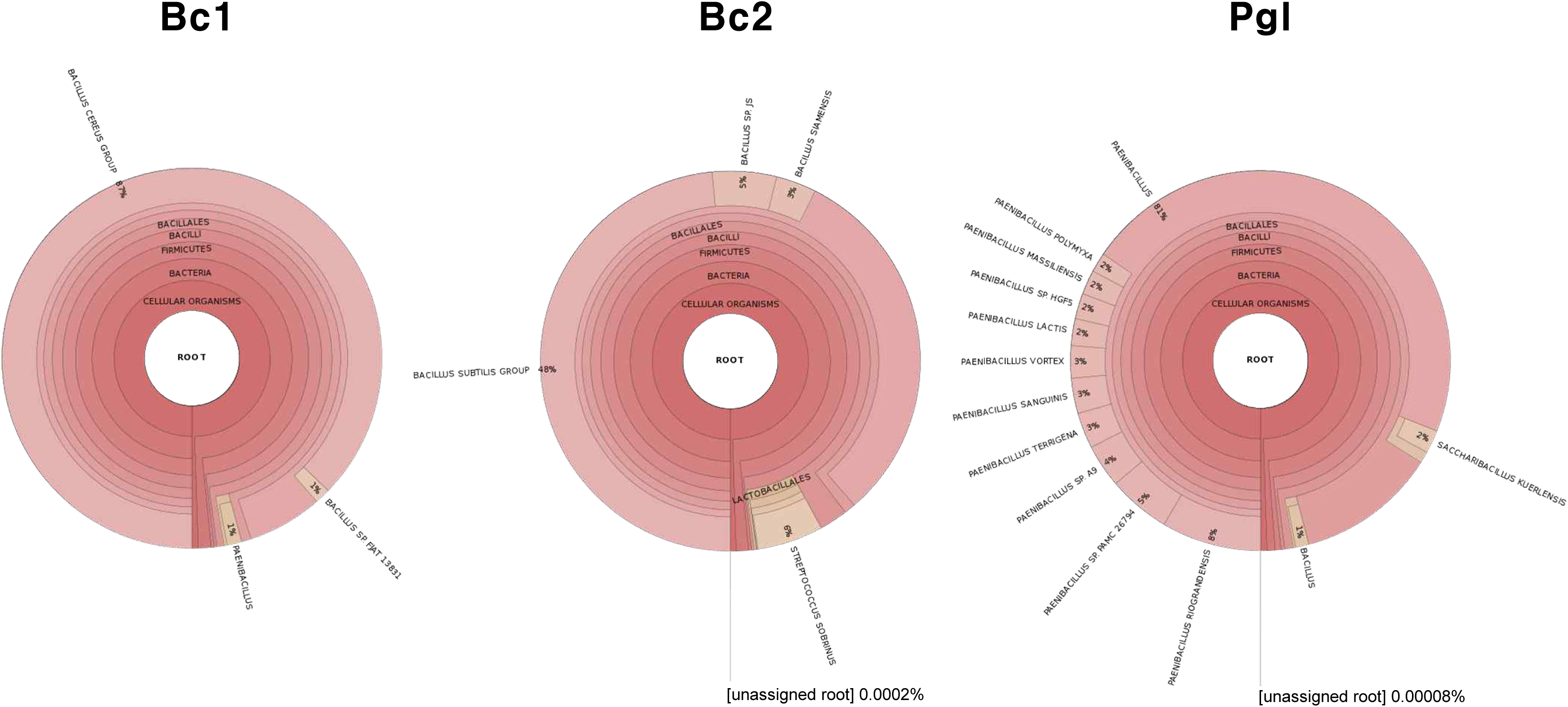
Krona plots of PhyloSift analysis results for selected samples Bc1, Bc2, and Pgl.

### *De novo* assemblies of all samples were significantly improved after k-mer abundance filtering

We previously showed that read filtration based on k-mer abundance could improve *de novo* assembly of samples Bc1 and Bp1 by removing less-abundant reads originating from contamination. Further comprehensive analyses revealed that nearly all samples had contamination at different levels. Except for Bc2 whose secondary peak in k-mer spectrum was not far enough from the main one to determine a suitable coverage cutoff, all twelve samples were filtered at k-mer abundance of 50 and subjected to *de novo* assembly. As shown in Fig. 7, the assemblies for all samples were greatly improved in terms of contig numbers. Because short and low-coverage contigs were not formed after filtering out those low-frequency reads, there was no significant loss in total contig length or N50 length.

**Fig. 7.**
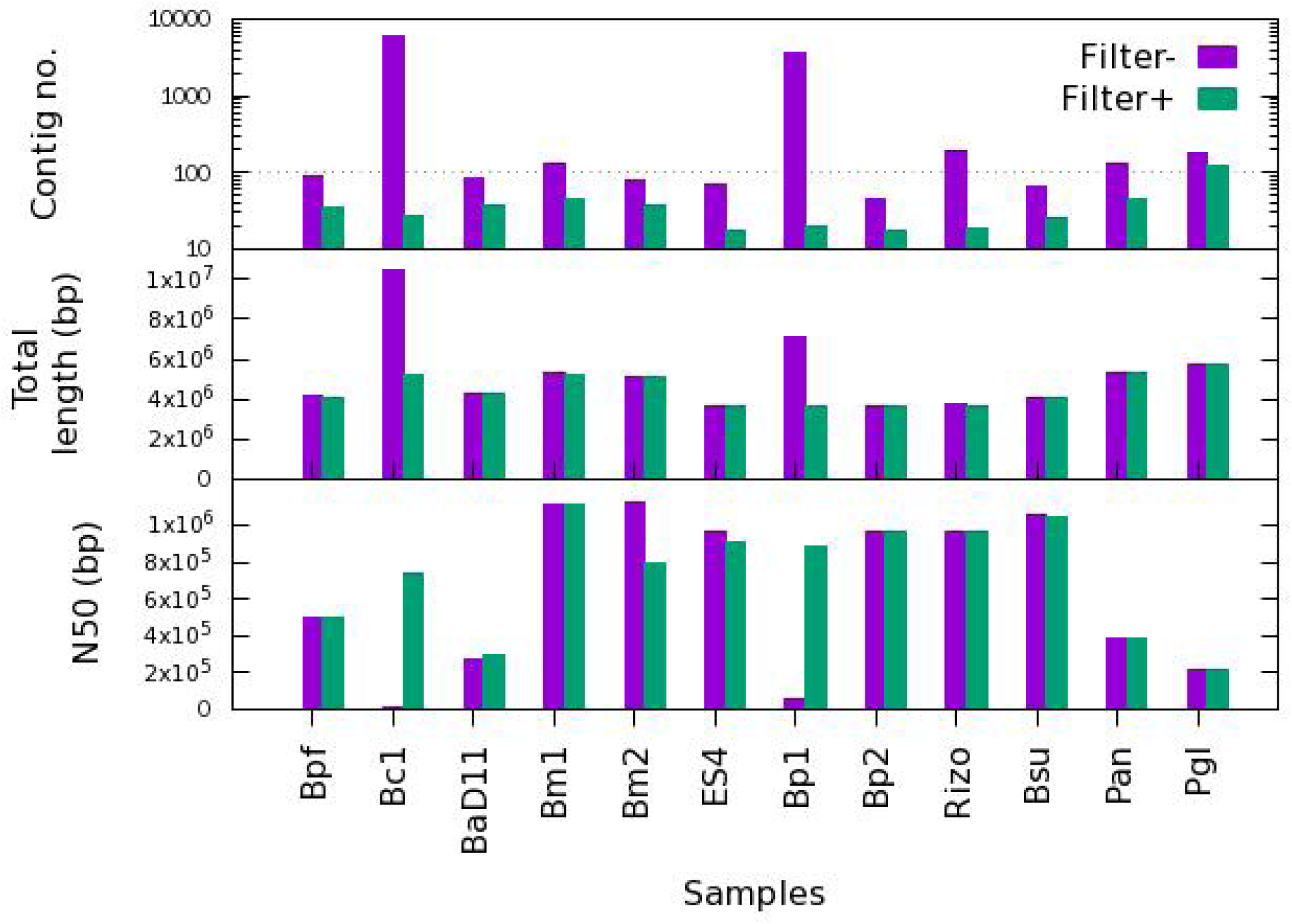
*De novo* assembly results after k-mer filtering of reads for twelve of the 13 samples (except Bc2).

## DISCUSSION

In this study, we took a systematic approach to determine the cause of failed Illumina assembly of bacterial genomes. Counting k-mers in sequencing reads with visualization was chosen as a quick and efficient method for the diagnosis of sequence reads. Sequencing error itself was not a crucial factor for successful *de novo* assembly as pretreatment such as trimming and error correction did not improve the results when sufficient sequencing depth, e.g., 100x, was ensured. Analyses using test datasets showed that the presence of repeats, contamination, and heterozygosity were the main reasons leading to poor Illumina assemblies for microbial genomes, which accompany minor peaks for k-mer abundance distribution.

At the first glance at assembly results, three (Bc1, Bc2, and Bp1) out of thirteen samples appeared to be problematic in terms of contig number and total contig length. In particular, Bc1 and Bp1 produced thousands of contigs whose total length were 3.9-5.0 Mb, which were larger than we expected. Coverage-length analysis showed that these contigs were divided into two groups based on average read coverage. The borderline of which was sequencing depth. The low-coverage contigs were usually short (< 1,000 kb). These short and low-coverage contigs comprised most of the contigs in the Bc1 and Bp1 assemblies. Re-mapping analysis indicated that reads responsible for the generation of short and low-coverage contigs ranged from 1.1% (Bp1) to 15.0% (Bc2) of total reads used. However, these numbers were negligible for other normal assemblies. This does not mean that contamination as low as ~1% of total reads is sufficient to spoil a *de novo* assembly. For example, only 1.9% of contaminating reads were mapped to short contigs when *E. coli* test dataset containing 5% *B. subtilis* reads were used. The remaining ~3% were unmapped, leaving only 3,347 reads (0.07% of total reads) mapped to high-coverage contigs.

K-mer spectra from real dataset leading to failed assemblies were very similar to those of test dataset with contamination. We could identify the source of contamination and estimate their proportions using several tools originally designed for metagenomics sequence analysis. Besides for specific contamination for individual samples (Bc1, Bc2, Bp1, Pan and Pgl), we unexpectedly found that all thirteen samples belonging to *Firmicutes* (*Bacillus* or *Paenibacillus* strains) had proteobacterial sequences. This does not imply true contamination across all samples, while the presence of caulobacterial reads were evident from our analysis results.

It is evident that contamination could cause difficulties in *de novo* assemblies or mislead conclusions. A tool for contamination removal called DeConSeq (34) has been introduced, but it cannot be used for unknown contaminants because it depends on alignment to known reference sequences. This tool is only useful for removing host sequences from metagenomics samples. Merchant *et al*. have reported cross-species contamination from public genome sequences (*Bos Taurus* and *Neisseria gonorrhoeae* TCDC-NG08107) using Kraken system (35). However, they did not suggest any possible contamination source other than erroneously labelled DNA. Recently, Mukherjee *et al*. have reported that PhiX sequence frequently used for a control during Illumina sequencing could be the source of contamination by screening publicly available microbial isolate genome sequences (36).

Then, what would be the cause of contamination in high-throughput genome sequencing experiments? In most cases, contamination could be due to insufficient axenic culture techniques of careless experimenters. Sometimes, bacterial stocks available from culture collections may be already contaminated. Simultaneous manipulation of multiple samples can also cause cross-contamination between samples. However, we suspect that the contamination with identical species, totally irrelevant to the laboratory environment, across multiple samples might have occurred during or after sequencing library preparation. Furthermore, bacterial cultivation and DNA preparation steps were undertaken at three independent laboratories in this study.

Insufficient wash between sequencing runs might cause carry-over contamination (http://seqanswers.com/forums/showthread.php?t=29110). In such a case, reads from a particular sample from the previous run could appear in the result of current sample whose barcode is coincident. In our results, however, multiple samples with different barcode showed similar patterns of contamination. It implies that contamination might have occurred during the library construction step at the sequencing facility. Very recently, we came across 1% archaeal reads and 6% eukaryotic reads from a myxobacterial sequencing result produced by the largest domestic sequencing company. Contamination caused by sequencing service provider poses a serious problem for the reliability of data, because it is very difficult for customers to take proper measures.

Not all contaminations are represented by anomalies in k-mer frequency distribution. We found that distinguishable k-mer frequency distribution could be hardly detected from simulated test dataset when contamination was 1% or less (data not shown). We cannot suggest any reference values for percent contamination that can ensure successful assembly. This can vary with actual situations. For example, 1% contamination can spoil assembly, while 5% contamination can generate reasonable assembly unless the contamination is dominated by one organism. However, there are a couple of things that we can consider, including diversity in contamination, sequence similarity between target organism and the contaminants, and sequencing quality. Even the choice of assembly program with a specific parameter set can greatly affect assembly results. Thus, as a general guideline, we propose to filter reads at a particular k-mer frequency (a quarter of sequencing depth would be suitable) for *de novo* assembly of microbial isolate genomes, even if contamination is not evident after the initial diagnosis using Jellyfish. Selecting contigs after the assembly step based on coverage would serve the same purpose. Read classification tools could then be applied for raw reads for deeper analysis of contamination, although they take long running time with the exception of MetaPhyler.

In summary, contamination is no longer a problem specific to individual samples in contemporary high-throughput sequencing era. Although not all incidence of contamination is reflected by unusual assembly metric values, a systematic analysis can reveal the underlying problem throughout samples. Our experience might provide insights on the utility of raw sequencing data for quality assessment of draft genome assemblies.

## ACKKNOWLEDGMENTS

We thank Tai-Boong Uhm (Chonbuk National University, Republic of Korea) and Yoav Bashan (The Bashan Institute of Science, USA) for providing bacterial strains for sequencing. We also thank Young Mi Sim (Korean Bioinformation Center) for technical assistance. This work was supported by KRIBB Research Initiative Program (KGM2111622) and the Bio & Medical Technology Development Program (NRF-2010-0029345) funded by Ministry of Science, ICT, and Future Planning, Republic of Korea.

## REFERENCES

1. Loman NJ, Pallen MJ. 2015. Twenty years of bacterial genome sequencing. Nat Rev Microbiol 13:787‐794.

2. Pop M, Salzberg SL. 2008. Bioinformatics challenges of new sequencing technology. Trends Genet 24:142‐149.

3. El-Metwally S, Hamza T, Zakaria M, Helmy M. 2013. Next-generation sequence assembly: four stages of data processing and computational challenges. PLoS Comput Biol 9:e1003345.

4. Birney E. 2011. Assemblies: the good, the bad, the ugly. Nat Methods 8:59‐60.

5. Junemann S, Prior K, Albersmeier A, Albaum S, Kalinowski J, Goesmann A, Stoye J, Harmsen D. 2014. GABenchToB: a genome assembly benchmark tuned on bacteria and benchtop sequencers. PLoS One 9:e107014.

6. Magoc T, Pabinger S, Canzar S, Liu X, Su Q, Puiu D, Tallon LJ, Salzberg SL. 2013. GAGE-B: an evaluation of genome assemblers for bacterial organisms. Bioinformatics 29:1718‐1725.

7. Koren S, Treangen TJ, Hill CM, Pop M, Phillippy AM. 2014. Automated ensemble assembly and validation of microbial genomes. BMC Bioinformatics 15:126.

8. Liao YC, Lin HH, Sabharwal A, Haase EM, Scannapieco FA. 2015. MyPro: a seamless pipeline for automated prokaryotic genome assembly and annotation. J Microbiol Methods 113:72‐74.

9. Koren S, Phillippy AM. 2015. One chromosome, one contig: complete microbial genomes from long-read sequencing and assembly. Curr Opin Microbiol 23:110‐120.

10. Chin CS, Alexander DH, Marks P, Klammer AA, Drake J, Heiner C, Clum A, Copeland A, Huddleston J, Eichler EE, Turner SW, Korlach J. 2013. Nonhybrid, finished microbial genome assemblies from long-read SMRT sequencing data. Nat Methods 10:563‐569.

11. Liao YC, Lin SH, Lin HH. 2015. Completing bacterial genome assemblies: strategy and performance comparisons. Sci Rep 5:8747.

12. Zerbino DR, Birney E. 2008. Velvet: algorithms for de novo short read assembly using de Bruijn graphs. Genome Res 18:821‐829.

13. Alkan C, Sajjadian S, Eichler EE. 2011. Limitations of next-generation genome sequence assembly. Nat Methods 8:61‐65.

14. Marcais G, Kingsford C. 2011. A fast, lock-free approach for efficient parallel counting of occurrences of k-mers. Bioinformatics 27:764‐770.

15. Anvar SY, Khachatryan L, Vermaat M, van Galen M, Pulyakhina I, Ariyurek Y, Kraaijeveld K, den Dunnen JT, de Knijff P, t Hoen PA, Laros JF. 2014. Determining the quality and complexity of next-generation sequencing data without a reference genome. Genome Biol 15:555.

16. Crusoe MR, Alameldin HF, Awad S, Boucher E, Caldwell A, Cartwright R, Charbonneau A, Constantinides B, Edvenson G, Fay S, Fenton J, Fenzl T, Fish J, Garcia-Gutierrez L, Garland P, Gluck J, Gonzalez I, Guermond S, Guo J, Gupta A, Herr JR, Howe A, Hyer A, Harpfer A, Irber L, Kidd R, Lin D, Lippi J, Mansour T, McA'Nulty P, McDonald E, Mizzi J, Murray KD, Nahum JR, Nanlohy K, Nederbragt AJ, Ortiz-Zuazaga H, Ory J, Pell J, Pepe-Ranney C, Russ ZN, Schwarz E, Scott C, Seaman J, Sievert S, Simpson J, Skennerton CT, Spencer J, Srinivasan R, Standage D, Stapleton JA, Steinman SR, Stein J, Taylor B, Trimble W, Wiencko HL, Wright M, Wyss B, Zhang Q, Zyme E, Brown CT. 2015. The khmer software package: enabling efficient nucleotide sequence analysis. F1000Res 4:900.

17. Jeong H, Kim HJ, Lee SJ. 2015. Complete genome sequence of Escherichia coli strain BL21. Genome Announc 3:e00134‐00115.

18. Jeong H, Sim YM, Park SH, Choi SK. 2015. Complete genome sequence of Bacillus subtilis strain ATCC 6051a, a potential host for high-level secretion of industrial enzymes. Genome Announc 3:e00532‐00515.

19. Jeong H, Lee DH, Ryu CM, Park SH. 2016. Toward complete bacterial genome sequencing through the combined use of multiple next-generation sequencing platforms. J Microbiol Biotechnol 26:207‐212.

20. Huang W, Li L, Myers JR, Marth GT. 2012. ART: a next-generation sequencing read simulator. Bioinformatics 28:593‐594.

21. Kim YS, Jeong JO, Cho SH, Jeong DY, Uhm T-B. 2012. Antimicrobial and biogenic aminedegrading activity of Bacillus licheniformis SCK B11 isolated from traditionally fermented red pepper paste. Kor J Microbiol 48:163‐170.

22. Bolger AM, Lohse M, Usadel B. 2014. Trimmomatic: a flexible trimmer for Illumina sequence data. Bioinformatics 30:2114‐2120.

23. Coil D, Jospin G, Darling AE. 2015. A5-miseq: an updated pipeline to assemble microbial genomes from Illumina MiSeq data. Bioinformatics 31:587‐589.

24. Simpson JT, Durbin R. 2010. Efficient construction of an assembly string graph using the FM-index. Bioinformatics 26:i367‐373.

25. Gurevich A, Saveliev V, Vyahhi N, Tesler G 2013. QUAST: quality assessment tool for genome assemblies. Bioinformatics 29:1072‐1075.

26. Seemann T. 2014. Prokka: rapid prokaryotic genome annotation. Bioinformatics 30:2068‐2069.

27. Mende DR, Sunagawa S, Zeller G, Bork P. 2013. Accurate and universal delineation of prokaryotic species. Nat Methods 10:881‐884.

28. Yuan C, Lei J, Cole J, Sun Y. 2015. Reconstructing 16S rRNA genes in metagenomic data. Bioinformatics 31:i35‐43.

29. Chun J, Lee JH, Jung Y, Kim M, Kim S, Kim BK, Lim YW. 2007. EzTaxon: a web-based tool for the identification of prokaryotes based on 16S ribosomal RNA gene sequences. Int J Syst Evol Microbiol 57:2259‐2261.

30. Liu B, Gibbons T, Ghodsi M, Treangen T, Pop M. 2011. Accurate and fast estimation of taxonomic profiles from metagenomic shotgun sequences.BMC Genomics 12 Suppl 2:S4.

31. Wood DE, Salzberg SL. 2014. Kraken: ultrafast metagenomic sequence classification using exact alignments. Genome Biol 15:R46.

32. Darling AE, Jospin G, Lowe E, Matsen FAt, Bik HM, Eisen JA. 2014. PhyloSift: phylogenetic analysis of genomes and metagenomes. PeerJ 2:e243.

33. Ondov BD, Bergman NH, Phillippy AM. 2011. Interactive metagenomic visualization in a Web browser. BMC Bioinformatics 12:385.

34. Schmieder R, Edwards R. 2011. Fast identification and removal of sequence contamination from genomic and metagenomic datasets. PLoS One 6:e17288.

35. Merchant S, Wood DE, Salzberg SL. 2014. Unexpected cross-species contamination in genome sequencing projects. PeerJ 2:e675.

36. Mukherjee S, Huntemann M, Ivanova N, Kyrpides NC, Pati A. 2015. Large-scale contamination of microbial isolate genomes by Illumina PhiX control. Stand Genomic Sci 10:18.

